# Defense-associated reverse transcriptases synthesize DNA in a telomerase-like manner

**DOI:** 10.64898/2026.10.08.757509

**Authors:** Shuai Chen, Shuqin Zhang, Youjing Zhang, Pengyu Fu, Tongyu Mao, Yancheng Zhan, Jie Han, Zhikun Liu, Zhengda Shan, Heng Zhang, Zhiyong Yuan, Ying Yu, Hang Yin

**Author notes:** These authors contribute equally (Shuai Chen, Shuqin Zhang, Youjing Zhang).

## Abstract

Defense-associated reverse transcriptases (DRTs) constitute an emerging class of bacterial defense systems that harness reverse-transcriptase activity to generate cDNA products and counter phage infection. A distinct subset of DRT10 systems evolutionarily related to telomerase generates tandem-repeat DNA to confer antiphage immunity. Here, we biochemically characterize the reverse transcription activity of DRT10 and determine cryo-EM structures of DRT10-ncRNA complex in three functional states, revealing an unanticipated 2:1 RT:ncRNA assembly mode. Two DRT10 protomers dimerize through a central pair of interlocked β-hairpins and are further embraced by a single ncRNA. This architecture distinguishes DRT10 from previously characterized DRT complexes, which either lack an ncRNA component or exhibit an equivalent stoichiometry of RT:ncRNA. Strikingly, despite the nearly identical conformations of the two DRT10 protomers, cDNA product density was resolved in only one catalytic pocket, revealing different product occupancy within the dimer. Moreover, along with flanking linkers, the stem loops of the ncRNA as fixed points to supply steric barriers help define the template boundary. These findings reveal a distinctive ncRNA-mediated DRT10 dimeric assembly and establish a mechanistic framework for protein-primed reverse transcription constrained by the length of the RNA flanking linker.

## Introduction

The evolutionary arms race between hosts and invaders expands the diverse arsenals of antiviral immune systems. Many well-characterized systems, including CRISPR-Cas systems, rely on the recognition and degradation of invasive nucleic acids, whereas emerging reverse-transcriptase-based systems use nucleic acid synthesis as a defense strategy (1–3). Reverse transcriptase activity was independently reported in 1970 (4,5), establishing the enzymatic basis for the transfer of genetic information from RNA to DNA. Beyond their canonical roles in retroelement propagation, reverse transcriptases (RTs) have increasingly been co-opted to mediate antiviral defense through diverse molecular mechanisms. For example, retrons are bacterial genetic elements employing an RT component to use non-coding RNA (ncRNA) template to generate multi-copy single-stranded DNA (msDNA). The RT-ncRNA-msDNA complex restrains toxic effectors until phage-induced disassembly of these complexes releases toxic effector activity (6). CRISPR-associated RTs enable the acquisition of RNA-derived spacers for immune memory storage (7).

Recent work has provided detailed mechanistic insights into several UG/Abi systems, revealing striking diversity in their modes of DNA synthesis and antiviral defense strategies. UG/Abi RTs are categorized into three classes (8). Among αRep-containing Class 1 UG/Abi RTs, the dual-RT DRT3 system coordinates ncRNA- and protein-templated DNA synthesis to generate dsDNA, which accumulates upon phage-mediated inhibition of RecBCD-dependent degradation and triggers abortive infection (9,10). DRT4-mediated protein-primed DNA synthesis promotes RNase activation, thereby resulting in cell growth arrest (11,12). Class 2 UG/Abi RTs, which contain an RT domain without additional annotated effector domains, employ different ncRNA-guided strategies. DRT2 uses rolling-circle reverse transcription to generate repetitive cDNA, which is subsequently transcribed and translated into the toxic repetitive Neo protein, resulting in growth arrest (2,3). Instead, DRT9 catalyzes protein-primed, ncRNA-templated synthesis of long poly(dA) products, whose accumulation triggers abortive infection (13,14). In DRT1 from Class 3 UG/Abi RTs, which is associated with a nitrilase or phosphohydrolase domain, template-independent DNA synthesis produces covalent protein-DNA adducts that promote assembly of a pre-active filament, priming the system for phage-triggered cell death (15). Collectively, these findings establish UG/Abi RTs as a mechanistically diverse class of immune systems that harness distinct modes of nucleic acid synthesis to deploy novel antiviral defense. Despite these advances, many UG/Abi RT systems remain mechanistically uncharacterized, highlighting the potential to reveal novel mechanisms of nucleic acid synthesis and antiviral defense. DRT10 is a recently identified defense-associated reverse transcriptase system evolutionarily related to telomerase and synthesizes tandem-repeat DNA (16).

Inspired by the elegant work, we characterize an *Escherichia coli* UG17 (DRT10) triple system, comprising the reverse transcriptase DRT10, an ncRNA and a predicted transmembrane protein, SLATT. We show that both the RT activity of DRT10 and SLATT are required for the antiphage defense. Biochemical and structural analyses reveal that DRT10 catalyzes protein-primed, ncRNA-templated cDNA synthesis. Cryo-EM structures of the DRT10-ncRNA complex in three functional states further reveal an ncRNA-mediated dimeric architecture and delineate a stepwise catalytic mechanism involving substrate-induced rearrangements, conserved protein-primed synthesis and stabilization of the emerging RNA-cDNA duplex. Together, our findings establish DRT10 as a distinct RT-based triple immune system and provide a mechanistic framework for understanding how a conformationally dynamic ncRNA coordinates complex assembly and cDNA synthesis.

## Results

### Biochemical characterization of the DRT10 system

To probe whether the DRT10 system confers antiphage defense, we investigated a DRT10 system derived from *Escherichia coli* H120 (**Figure 1a**). The DRT10 system consists of a DRT10 protein containing a predicted reverse transcriptase (RT) domain, a non-coding RNA (ncRNA) and a SLATT protein containing a predicted transmembrane region. To identify the components essential for DRT10-mediated antiphage defense, we challenged MG1655 cells carrying an empty vector or the wild-type or mutant DRT10 system with serial tenfold dilutions of phage Bas18 (**Figure 1b**). Specifically, plaque assays showed that either deletion of SLATT component or catalytic inactivation of the DRT10 protein (D232A/D233A) abolished antiphage defense, indicating that both the SLATT protein and the reverse transcriptase activity of the DRT10 protein are required for the DRT10 system to execute antiphage activity. This distinguishes the DRT10 system from the previously characterized DRT2, DRT3 and DRT9 defense systems (2,3,9,10,13,14), which perform defense activity in the absence of other ancillary components, implying mechanistic diversity among DRT systems.

**Figure 1.**
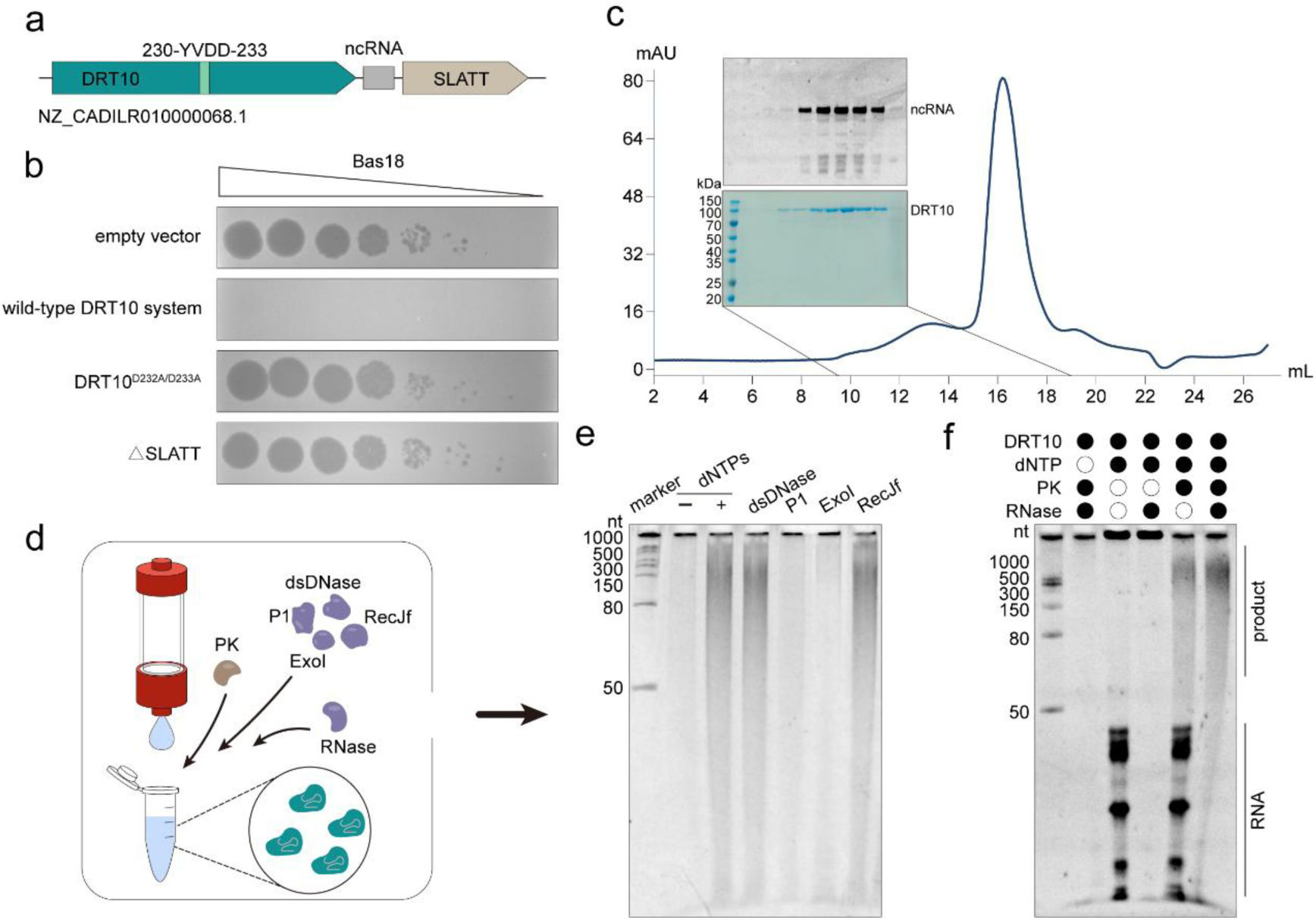
Biochemical characterizations of the DRT10 antiphage system. **a** Gene architecture of the DRT10 system from *Escherichia coli* H120. The catalytic residues of the DRT10 protein are indicated. **b** Plaque assay with serial dilution of phage Bas18 spotted onto lawns of MG1655 cells harboring an empty vector or plasmids encoding wild-type or mutant DRT10 systems. Images are representative of three independent biological replicates. **c** Purification of the DRT10-ncRNA complex using size exclusion chromatography. The resulting peak fractions were analyzed using urea-PAGE and SDS-PAGE. **d** Schematic workflow for the identification of DRT10-generated reverse-transcription products. The purified DRT10-ncRNA complex was incubated with dNTPs and subsequently treated with proteinase K (PK). Following heat inactivation of proteinase K, aliquots were separately incubated with dsDNase, P1, ExoI or RecJf. After RNase treatment, the samples were analyzed by urea-PAGE. **e** Urea-PAGE analysis of the reverse transcription product subjected to the indicated treatments. **f** Urea-PAGE analysis of products synthesized by DRT10. Gel images are representative of three independent experiments.

To further assess the reverse transcriptase activity of DRT10, we purified the DRT10-ncRNA complex and conducted the reverse transcription assay in vitro (**Figure 1c, Figure S1**). Product accumulation was observed with increasing incubation time, dNTP concentrations and salt concentrations (**Figure S1**). Furthermore, the cDNA products were degraded by ExoI, a 3’-to-5’ single-stranded DNA exonuclease, but resisted RecJf, which degrades single-stranded DNA in the 5’ to 3’ direction (**Figure 1d-e**). Moreover, discrete cDNA bands became detectable only after protease digestion, indicating the release of protein-linked reverse-transcription products (**Figure 1f**). Collectively, these findings support a protein-primed mechanism in which DRT10 remains covalently attached to the 5’ end of the nascent cDNA. This resembles those of DRT9, AbiA and AbiK (13,14,17,18), suggesting that protein-primed cDNA synthesis is a conserved mechanism among members of the UG/Abi RT family.

### Architecture of the DRT10 system in the dNTP-free state

To gain further insight into the structural basis of the DRT10 system, we determined the structure of the DRT10-ncRNA binary complex in the absence of dNTPs at a high resolution of 2.59 Å (termed dNTPs-free state hereinafter) (**Figure S2**). The high-quality density map enabled us to build the atomic model for most of the DRT10 protein and the ncRNA, with the exception of the carboxyl-terminal loop of DRT10 (residues 466-469) and a six-nucleotide segment of the ncRNA (80U-85U), likely owing to their intrinsic conformational flexibility in the dNTP-free state.

Unanticipatedly, the DRT10-nRNA binary complex comprises two DRT10 protomers and a non-coding RNA molecule, adopting an overall pseudo-symmetric dimeric architecture (**Figure 2a**). The two DRT10 protomers assemble in a face-to-face configuration, with one protomer rotated by approximately 180° relative to the other (**Figure S3a-b**). The ncRNA threads through the assembly and extensively contacts the two protomers. DRT10 protomer adopts a right-hand-like architecture and comprises N-terminal extension (NTE, residues 1-40), fingers domain (residues 41-131 and 148-198), palm domain (residues 132-147 and 199-292), thumb domain (residues 293-384), and C-terminal extension (CTE, residues 385-469) (**Figure 2b-c**). Two DRT10 protomers adopt nearly identical conformations (RMSD = 0.623 Å) and associate mainly through a pair of interlocked β-hairpins (**Figure 2d, Figure S3b**).

**Figure 2.**
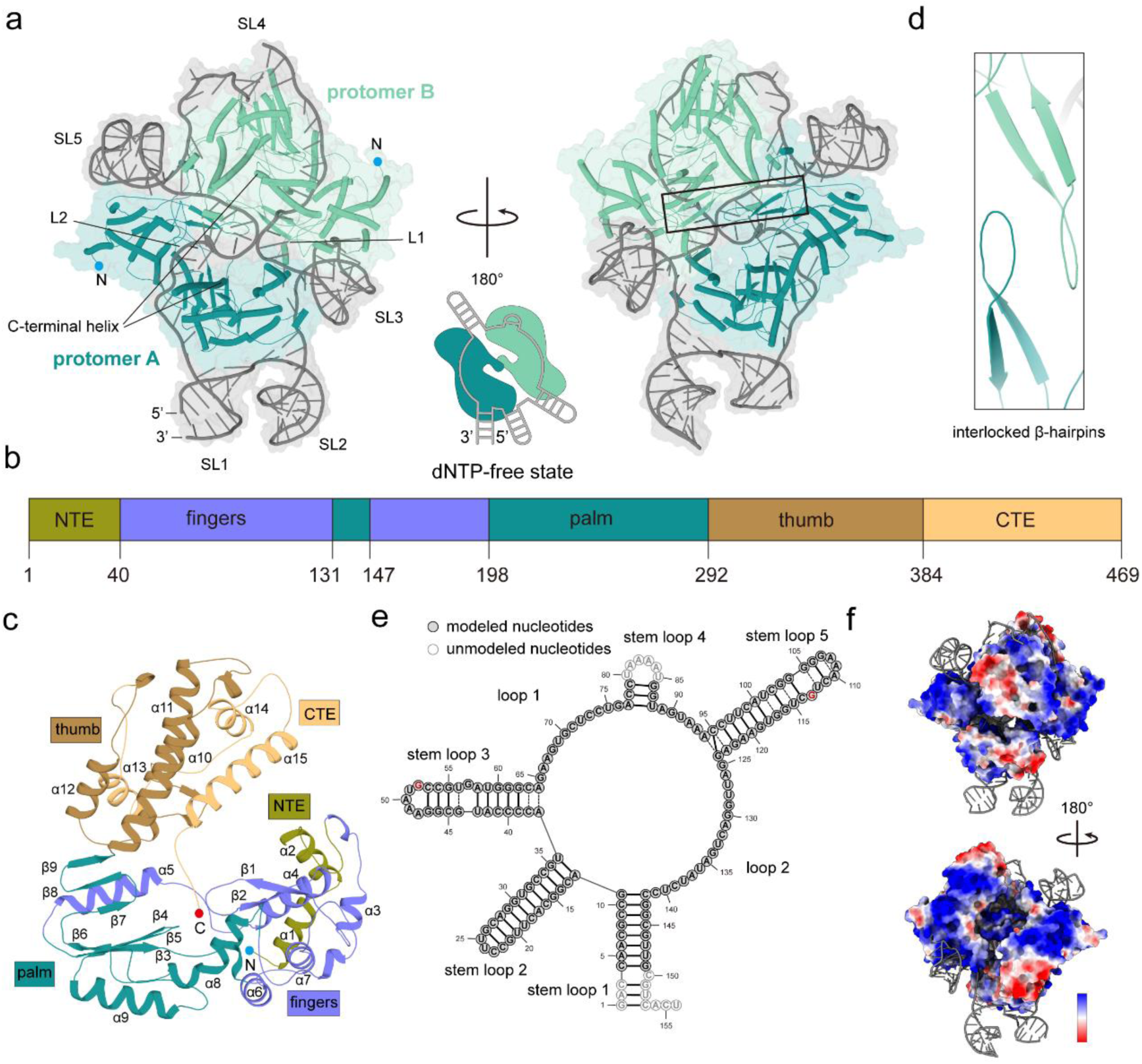
Overall structure of DRT10-ncRNA complex in the dNTP-free state. **a** Atomic model of DRT10-ncRNA complex in the dNTP-free state. A schematic representation of the complex assembly is shown. **b** Schematic representation of the domain organization of DRT10 protomer. **c** Structure of a DRT10 protomer, colored according to the domain organization shown in (**b**). **d** Close-up view of the interlocked β-hairpins at the DRT10 dimerization interface. **e** Schematic representation of the secondary structure of the DRT10-associated ncRNA. The modeled and unmodeled nucleotides are indicated. Nucleotides shown in red flip out and dock into pockets in the respective DRT10 protomers. **f** Electrostatic surface potential of the DRT10 dimer.

Protein-protein interactions in Class 1 UG/Abi RTs (e.g., AbiK and Abi-P2) and Class 2 UG/Abi RTs (e.g., DRT9) are mainly involved in helix-helix contacts (13,14,17). By contrast, a pair of interlocked β-hairpins forms the principal protein-protein interface within the DRT10 dimer. Specifically, the long β1-β2 β-hairpins protruding from the two protomers interlock in an antiparallel arrangement across the dimer interface (**Figure S3c**). Arg77, situated at the tip of the β-hairpin, inserts into a groove on the opposing protomer lined by Glu70, Leu72, Val83 and Thr178 from the adjacent α7-α8 loop. The sidechain of Arg77 hydrogen bonds with Glu70 and Thr178. Deletion of residues 76-79 impaired both reverse-transcription activity and antiphage defense, supporting an important role for the interlocked β-hairpins in DRT10 function (**Figure S3d-e**). On the opposite side of the DRT10 dimer, ncRNA is embedded within a large cleft formed by the fingers, thumb and CTE domains of the corresponding DRT10 protomer, thereby positioning unpaired RNA within the catalytic pockets (**Figure 2**).

The ncRNA folds into four extended stem loops (stem loops 1-3 and stem loop 5), one compact stem loop 4, and two long loops (loop 1 and loop 2) (**Figure 2e**). The stem loops collectively form a compact scaffold that embraces the DRT10 dimer, whereas loop 1 and loop 2 traverse the respective catalytic pockets of the two DRT10 protomers. The DRT2-ncRNA complex functions as a monomer during antiphage defense, whereas the DRT9-ncRNA complex assembles into a hexamer in which each ncRNA engages an adjacent DRT9 protomer and forms a kissing-loop interaction with a neighboring ncRNA (3,13,14). Unlike these characterized ncRNA-dependent DRT systems, the ncRNA in the DRT10 system spans both DRT10 protomers and contributes to the dimeric architecture (**Figure 2f**). Stem loop 1 and stem loop 2 protrude outwards away from the complex, whereas stem loop 3 and stem loop 5 dock onto the fingers domains of the respective DRT10 protomers, establishing extensive protein-RNA contacts (**Figure S4a-b**). A patch of positively charged residues engages both the major and minor grooves of these stem loops, forming electrostatic contacts and hydrogen bonds with their phosphate-ribose backbones. Interestingly, both 52G and 113G are well resolved in the density map, with each nucleotide accommodated in an equivalent pocket lined by Arg64, Phe65, Arg172, Leu175 and Thr176. The guanine base forms hydrogen bonds with the backbone carbonyl oxygen of Phe65 and the side chain of Arg172, while being underpinned by Leu175 and Thr176. Additionally, the ncRNA also contains several noncanonical base pairs, including G-U, A-A and C-C pairs (**Figure 2e**). The side chains of Arg37 and Arg380 extend towards the minor and major grooves, respectively, and contribute to the hydrogen-bonding networks surrounding these noncanonical pairs, further stabilizing stem loop 3 and stem loop 5 (**Figure S4c**). Together, these interactions fix stem loop 3 and stem loop 5 to the DRT10 dimer and support their local folds.

### Substrate-induced rearrangements in the dNTP-bound state

To elucidate the molecular basis of DRT10-mediated reverse transcription, we incubated the DRT10-ncRNA complex with dNTPs and determined cryo-EM structures in the dNTP-bound and product-bound states at overall resolutions of 2.81 and 3.44 Å, respectively (**Figure 3, Figure S5**). Clear densities corresponding to bound dNTPs and nascent cDNA product were observed in the final reconstructions (**Figure 4, Figure S6**). Structural comparison between the dNTP-free and dNTP-bound states reveals no substantial changes in the overall domain arrangement of the complex (**Figure S6a**). Nevertheless, dNTP binding and cDNA synthesis were accompanied by localized rearrangements of key structural elements surrounding the catalytic pockets and the RNA template regions (discussed below).

**Figure 3.**
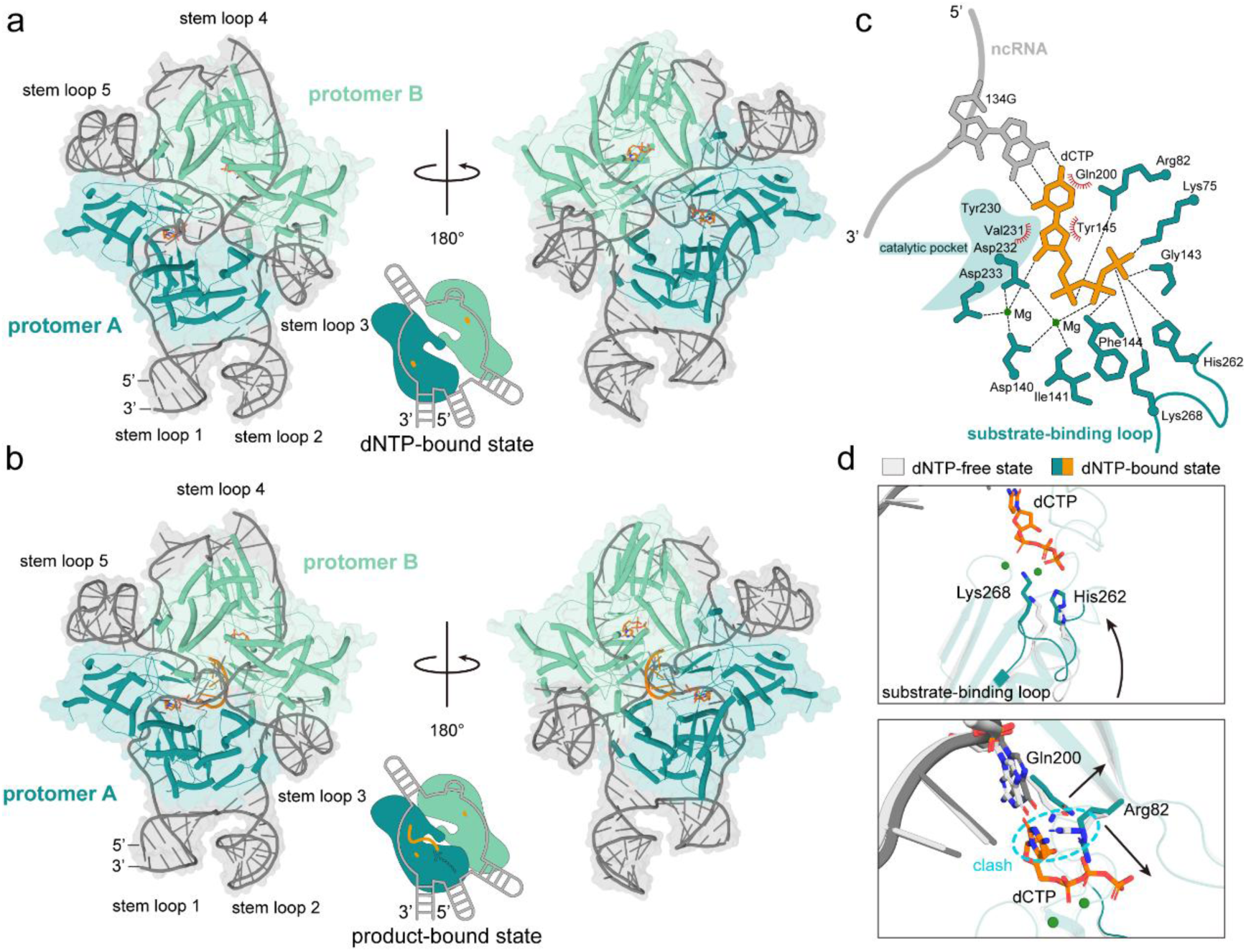
A key structural element is responsible for substrate engagement in DRT10. **a-b** Overall structures of DRT10-ncRNA complex in the dNTP-bound and product-bound states. **c** Schematic representation of the interactions coordinating the incoming dNTP within the DRT10 catalytic pocket. The shaded region indicates the conserved catalytic motif. **d** Structural comparison of the DRT10-ncRNA complex in the dNTP-free and dNTP-bound states. Key residues are shown as sticks and conformational changes are indicated by arrows. The blue circle highlights a potential steric clash.

**Figure 4.**
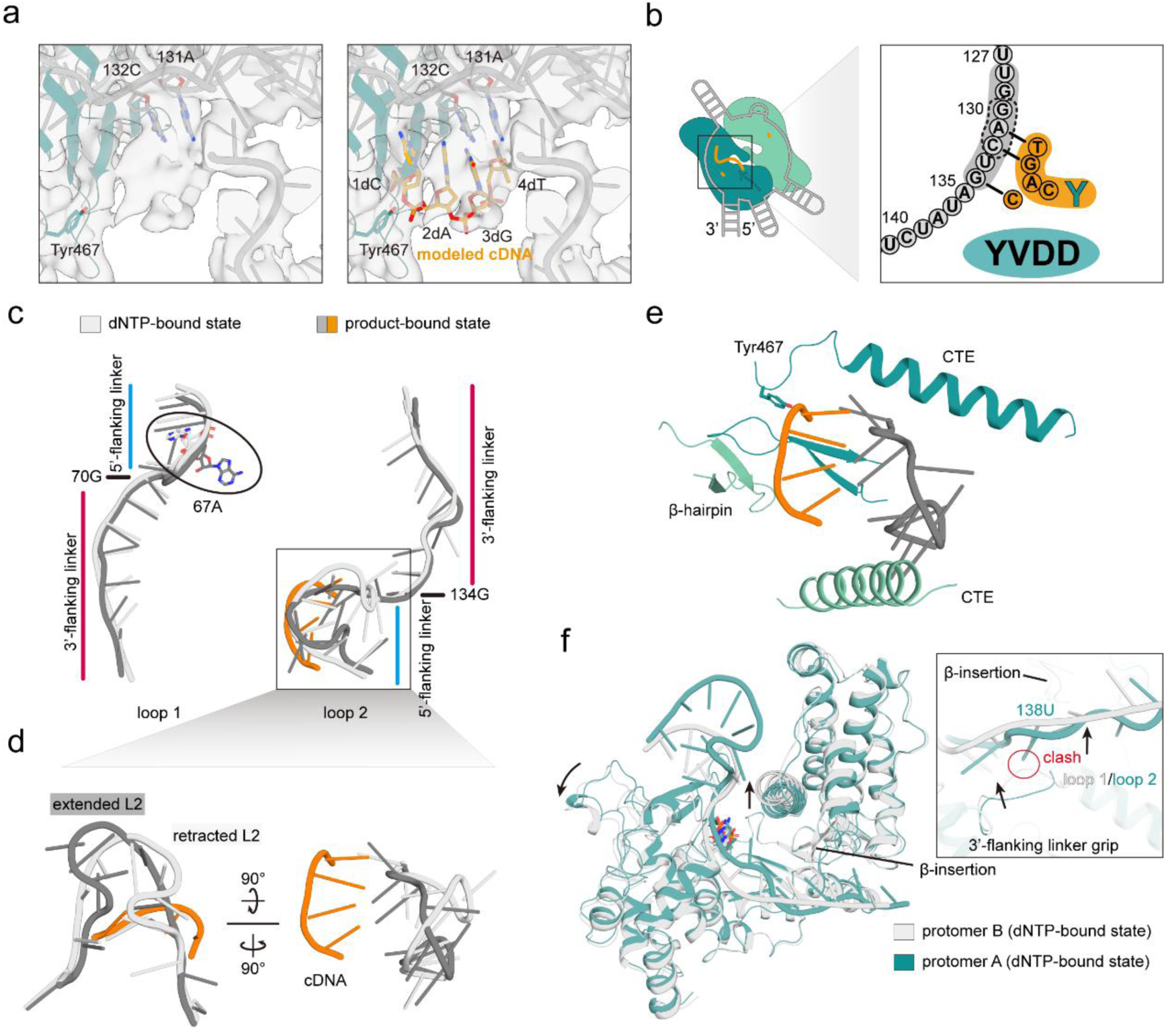
ncRNA guides the product synthesis in the DRT10-ncRNA complex. **a** Close-up view of the cryo-EM density map corresponding to the synthesized cDNA in the product-bound state. **b** Schematic representation of the DRT10-ncRNA complex. The enlarged view on the right shows the nascent cDNA product and its base-pairing interactions with the ncRNA template. **c** Structural comparison of loop 1 (left) and loop 2 (right) in the dNTP-bound and product-bound states. The flipped nucleotide 67A is highlighted by a black oval and the product-template base-paired region is indicated by a black box. **d** Detailed view showing the structural rearrangement of the 5’-flanking linkers in loop 2. **e** The product-template base pairs are supported from below by the interlocked β-hairpins and flanked by the two CTEs. **f** Structural comparison of DRT10 protomers A and B in the dNTP-bound state. Zoomed view represents the structural rearrangements of the 3’-flanking linker grip and 3’-flanking linkers in loop 1 and loop 2 are indicated by arrows. Potential steric clashes are highlighted by red circles.

We first examined the dNTP-bound state to investigate how incoming dNTPs are accommodated within the DRT10 catalytic pockets. In the dNTP-bound state, well-defined densities for the incoming dNTPs are observed within each catalytic pocket of DRT10, adjacent to the conserved YXDD catalytic motif (**Figure S6b**). The substrate is stabilized by an extensive network of metal-coordinated and protein-mediated interactions (**Figure 3c**). The sidechains of Asp140, Asp232 and Asp233, together with the backbone carbonyl oxygen of Ile141, coordinate two Mg^2+^ ions, which also engage the triphosphate moiety of the incoming dCTP. His262 forms a hydrogen bond with the γ-phosphate. Lys75 and Lys268, lining the nucleotide-binding pocket, contact the negatively charged triphosphate group. Arg82 further engages the incoming dCTP by contacting the α-phosphates while stacking against the cytosine base. Phe144 and Tyr145 pack against the deoxyribose moiety of dCTP, further stabilizing the bound dCTP. The nucleotides 70G and 134G each form a Watson-Crick base pair with the cytosine base of the incoming dCTPs in the corresponding catalytic pocket. Together, these conserved residues engage the incoming dNTPs and position them for incorporation into the nascent cDNA (**Figure S6c**).

Structural comparison between the dNTP-free and dNTP-bound states reveals conformational changes induced by dNTP binding. Although the overall dimeric architecture remains unchanged (RMSD = 0.6 Å), several structural elements surrounding the catalytic pockets undergo notable rearrangements, especially within the palm domains (**Figure S6a**). In particular, a loop (aa 261-270, termed substrate-binding loop) shifts inward by approximately 3 Å towards the substrate-binding pocket, accompanied by a displacement of His262 and Lys268 (**Figure 3d**). This rearrangement positions His262 and Lys268 to engage the substrate and facilitate substrate stabilization in the dNTP-bound state. A comparable substrate-associated conformational change is also observed in DRT9 protein (**Figure S6d**). By contrast, no equivalent rearrangement was resolved in DRT2, possibly because its nucleotide-binding pocket is preorganized for substrate accommodation or because the corresponding conformational state is not captured. Moreover, upon substrate binding, the side chains of Arg82 and Gln200 reorient to avoid potential steric clash and stabilize the incoming dNTP within the catalytic pocket (**Figure 3d**). Notably, these substrate-interacting residues are conserved across DRT10 homologs, suggesting that this mode of dNTP accommodation and stabilization may be conserved within the DRT10 family (**Figure S6c**).

Collectively, the rearrangements triggered by dNTP binding facilitate the accommodation of the incoming dNTPs opposite the template nucleotides and position them for subsequent incorporation.

### Reverse transcription synthesis mechanism of the DRT10 system

Consistent with our biochemical evidence that the reverse transcription products were not processed by 5’-to-3’ DNA exonuclease (**Figure 1**), we unambiguously observe continuous density extending from Tyr467 in the C-terminal loop in the product-bound state (**Figure 4a**). This density allows us to model a four-nucleotide cDNA fragment (5’-CAGT-3’), in which 3dG and 4dT form Watson-Crick base pairs with the template nucleotides 132C and 131A, respectively (**Figure 4b**). In agreement with the structural evidence, the Y467F mutation disrupted the ability to synthesize the cDNA product and compromised antiphage defense (**Figure S3d-e, Figure S6e**). These biochemical and structural data support a Tyr467-mediated protein-primed mechanism for DRT10 reverse transcription, analogous to that reported for UG/Abi systems (13,14,17).

In comparison to the dNTP-bound state, the DRT10-ncRNA complex adopts a more compact dimeric architecture in the product-bound state (**Figure S7a-b**). The thumb and CTE domains undergo substantial rearrangements, accompanied by pronounced shifts of the associated RNA stem loops 1 and 2. Although loop 1 remains largely unchanged except for the base flipping of 67A, loop 2 adopts an extended conformation, contrasting with its relatively retracted configuration in the dNTP-bound state (**Figure 4c-d**). In this extended conformation, 131A and 132C flip towards the nascent cDNA and are positioned opposite their complementary bases, enabling the formation of product-template base pairs. The resulting RNA-cDNA duplex is further stabilized by the interlocked β-hairpin pair and the two CTE domains of the DRT10 dimer (**Figure 4e**). Few base-specific contacts are observed between DRT10 and either loop 1 or loop 2. Instead, most interactions involve the RNA phosphate backbone. The 5’ ends of loop 1 and loop 2 are fixed by stem loop 3 and stem loop 5, respectively (**Figure 2, Figure S4**). Their 3’ ends are constrained by stem loop 4 and stem loop 1, respectively. These structural elements fix the endpoints of loop 1 and loop 2 while allowing their intervening regions to adopt distinct conformations (**Figure 4**).

To understand how DRT10 defines the template region within the dimeric assembly, we take a close examination of the structures in the dNTP-bound and product-bound states (**Figure 4**). Particularly, we noticed that the 3’ flanking linkers of loop 1 and loop 2 contain six and seven nucleotides, respectively (**Figure 2e**). An RNA loop spanning residues 109-115 (termed 3’-flanking linker grip) adopts distinct conformations in two protomers. In comparison with DRT10 protomer B, reorientation of the 3’-flanking linker grip in protomer A is accompanied by outward shifts of the CTE and thumb domain, slightly elevating loop 1 and restricting the geometry of its 3’-flanking linker, shortening it by one nucleotide relative to the corresponding linker in loop 2 (**Figure 4f**). Compared with DRT2 and DRT9, DRT10 contains a distinctive β-insertion (residues 344-353) within the thumb domain that contacts the 3’-flanking linker and shifts together with the surrounding structural elements, suggesting a potential role in template definition. With stem loop 1 and stem loop 4 serving as steric barriers, the 3’-flanking linker and the 3’-flanking linker grip may therefore act together to delineate the 3’ template boundary and select the first nucleotide to be copied during reverse transcription.

At the 5’ ends of 5’-flanking linkers, His313 from each thumb domain stacks against the nucleotides 65G of loop 1 and 125G of loop 2, respectively (**Figure S7c**). The 5’-flanking linker of loop 1 contains five nucleotides, whereas its counterpart in loop 2 comprises nine, leading to a larger bulge in loop 2. A wedge loop spanning NTE and fingers domains (residues 28-46) supports both the flanking linkers and raises them above the protein surface. Notably, the conformations of the wedge loops and surrounding regions in protomers A and B are largely unchanged, implying that the length of the 5’-flanking linkers, rather than conformational changes, contributes to the different product occupancy observed in the product-bound state. Consistent with this possibility, loop 2 transitions from a retracted configuration in the dNTP-bound state to an extended configuration in the product-bound state (**Figure 4d**). This transition is reminiscent of the RNA accordion model proposed for telomerase, in which the 5’-flanking linker expands during cDNA synthesis and may subsequently return to its compact configuration under constraints imposed by its fixed boundaries (19–21) (**Figure 4d, Figure S7d-e**). Under these structural constraints, the wedge loop appears to help set the lower limit on the 5’-flanking linker length and determine the 5’ boundary of the template region. Thus, DRT10 combines fixed RNA stem loops with dynamic flanking linkers, allowing the protein and RNA scaffolds to help define the template boundaries while accommodating the structural changes associated with cDNA synthesis.

Together, we elucidate a stepwise mechanism in which dNTP binding reorganizes the catalytic pocket, Tyr467 primes cDNA synthesis, and subsequent conformational changes stabilize the emerging RNA-cDNA duplex.

## Discussion

Reverse transcriptases expand the routes of genetic information transfer by copying RNA into DNA. Although best known for their association with retroviruses, retrotransposons and telomere maintenance (22–25), there is emerging evidence that reverse transcriptases function as versatile molecular modules that support diverse biological processes, including antiviral immunity (6,8,26). Here, we establish DRT10 as an ncRNA-dependent reverse transcriptase that confers antiphage defense through a mechanism requiring both its catalytic activity and the predicted membrane-associated protein SLATT (**Figure 1**). Cryo-EM structures of the DRT10-ncRNA complex in the dNTP-free, dNTP-bound and product-bound states further reveal the molecular basis of ncRNA-mediated dimerization, substrate-associated conformational rearrangements and protein-primed cDNA synthesis (**Figure 5**).

**Figure 5.**
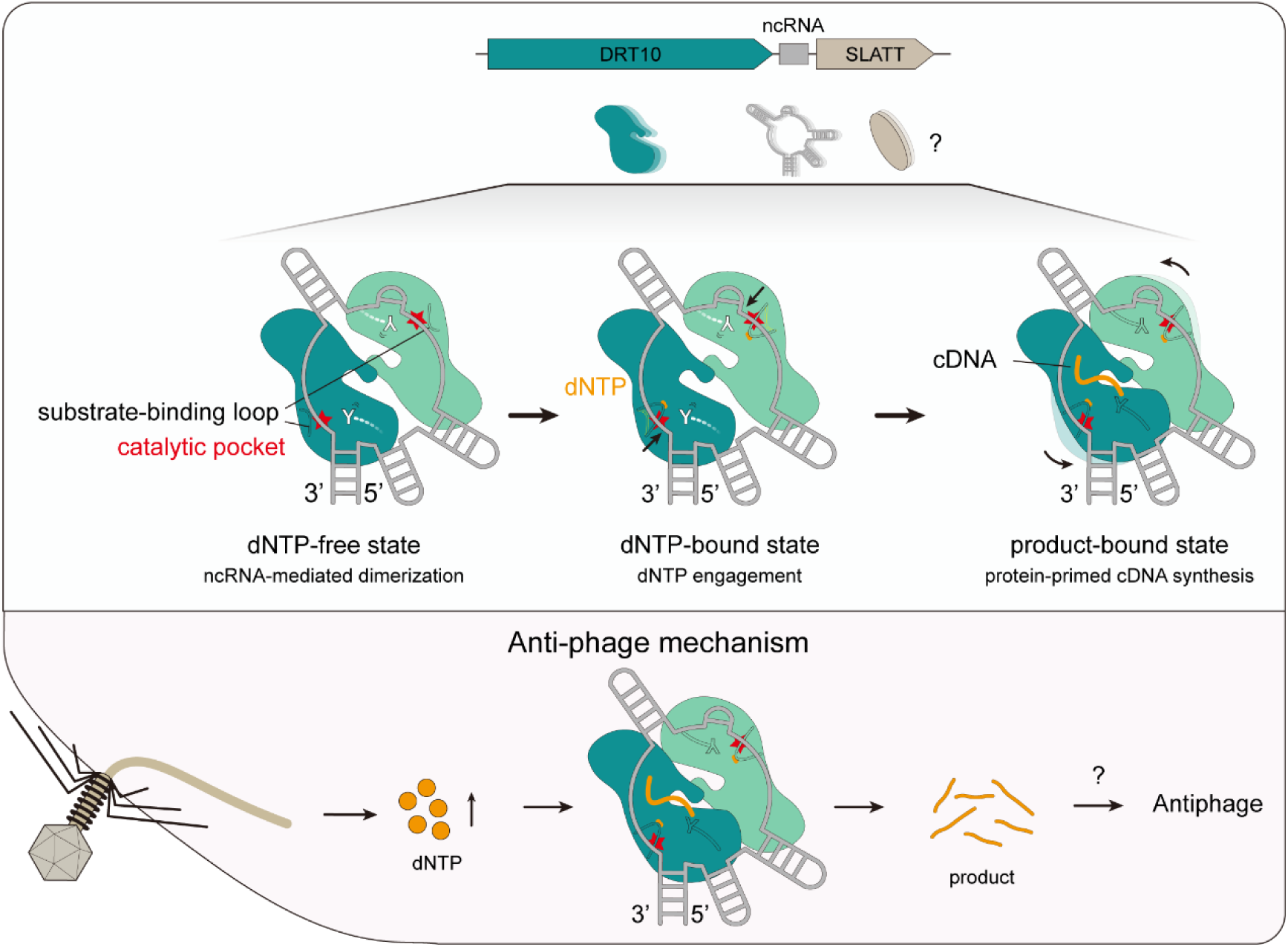
Proposed mechanism of the DRT10 system. In the tripartite DRT10 system, the DRT10 protein catalyzes protein-primed and template-dependent cDNA synthesis. Upon substrate binding, the substrate-binding loop rearranges to engage the incoming dNTP. As one DRT10 protomer synthesizes cDNA using the template in loop 2, the other protomer helps stabilize the nascent cDNA, thereby activating the antiphage defense.

Distinct from the previously characterized DRT systems in which a single RT protein or dual RT proteins are sufficient to confer antiphage defense (2,3,9–13,15), the DRT10 system requires both DRT10 and SLATT proteins to impart immunity. Disruption of either the DRT10 catalytic sites or SLATT abolished defense (**Figure 1**), indicating the functional relationship between cDNA synthesis and the membrane-associated component in the antiviral immune process. This highlights mechanistic diversification among the UG/Abi systems. Interestingly, in the DRT10-ncRNA complex, ncRNA, as a dual-functional element, serves as a structural scaffold to support the dimeric assembly while also providing the template to guide cDNA synthesis. Additionally, a pair of interlocked β-hairpins forms a distinctive dimerization interface, highlighting the diversity of assembly mechanisms among DRT systems. Moreover, we captured three structural snapshots to support a stepwise mechanism in which a functionally substrate-binding loop moves towards the catalytic pocket to engage the incoming dNTP. Subsequent coordinated rearrangements of the protein and RNA accommodate and stabilize the emerging RNA-cDNA duplex. The continuous density linking Tyr467 to the nascent cDNA, together with our biochemical evidence for protein-linked products, further supports protein-primed reverse transcription. This mechanism resembles that of DRT9, suggesting a prevalent protein-primed cDNA synthesis mechanism within this UG/Abi RT lineage. However, differences in oligomeric organization, template definition and accessory-protein dependence suggest considerable mechanistic diversity within the UG/Abi RTs.

The human telomerase can dimerize through its non-catalytic H/ACA RNP regions, without direct association between the two catalytic cores (27). Both catalytic pockets of dimeric human telomerase engaged DNA, whereas only one DRT10 catalytic pocket showed resolved density for the cDNA product. The different product occupancy observed in two structurally identical protomers suggests that conformational changes of reverse transcriptase are unlikely to be the primary determinant of this observation, implying a mechanism distinct from that of human telomerase. We propose that both catalytic pockets may be competent for cDNA synthesis, but differences in their 5’-flanking linkers could impose distinct constraints on template usage and, consequently, on the sequence and length of the cDNA produced. Nevertheless, the structures captured here do not distinguish whether only one pocket is catalytically active at a given time or whether the product-bound state in the second pocket is too transient or heterogeneous to be resolved. Notably, when this manuscript was in preparation, two independent studies reported similar findings on DRT10 (28,29). Their insightful studies elucidated template definition combining multiple mutations and the activation mechanism of the DRT10 system, respectively. Complementing these studies, our findings further reveal the mechanistic insights that cDNA is generated under the constraint of ncRNA and stabilized by the dimeric architecture, expanding the mechanistic repertoire of RT-mediated bacterial immunity (**Figure 5**).

## Methods

### Protein expression and purification

The coding region of full-length DRT10 protein and its associated ncRNA (Accession: NZ_CADILR010000068.1) were amplified by PCR and inserted into the 2CT-10 vector (Addgene #55209) with an N-terminal His10-MBP tag. The constructed plasmid was transformed into *E. coli* BL21-AI cells. When the cultures reached an OD_600_ of 0.8, 0.2 mM IPTG was supplemented to induce protein expression. After collection by centrifugation, cells were lysed by sonication in lysis buffer (25 mM Tris-HCl pH 7.5, 500 mM NaCl, 2 mM β-mercaptoethanol). The lysate was clarified by centrifugation at 14,000 rpm for 45 min at 4 °C, then purified by Ni-NTA affinity purification (Qiagen). The target complex was eluted with lysis buffer containing 300 mM imidazole. The resulting elution containing the tag-fused target complex was further purified by ion-exchange chromatography using a HiTrap Q HP column (Cytiva) and size-exclusion chromatography using a Superose 6 Increase column (Cytiva) equilibrated with gel-filtration buffer (25 mM Tris-HCl pH 7.5, 300 mM NaCl, 2 mM DTT). Peak fractions were analyzed by urea-PAGE and SDS-PAGE. The mutant DRT10-ncRNA complexes were expressed and purified as described above.

### In vitro reverse transcription assay

The purified protein complex was incubated with 500 μM dNTPs in a 20 μL reaction mixture containing 25 mM Tris-HCl pH 7.5, 500 mM NaCl, 5 mM MgCl_2_ and 2 mM DTT. After incubation at 37 °C for 1 h, the reaction was terminated by proteinase K for 30 min. Proteinase K was subsequently inactivated at 95 °C for 3 min, followed by RNase A digestion at 37 °C for 15 min. The resulting products were resolved by 15% urea-PAGE and visualized using an Azure Imaging System. The assays of the purified DRT10-ncRNA complex incubated with dNTPs were performed similarly over the increasing incubation times, dNTP concentrations and salt concentrations.

### SDS-PAGE analysis

To examine the covalent linkage of nucleic acid to DRT10 protein, the purified complex was incubated with increasing dNTPs or 500 μM dNTPs for 1 h at 37 °C in the reaction buffer (25 mM Tris-HCl pH 7.5, 500 mM NaCl, 5 mM MgCl_2_, 2 mM DTT). The reaction was terminated by RNase A, followed by heat denaturation at 95 °C for 3 min. Samples were separated by 12% SDS-PAGE and detected by Coomassie Brilliant Blue staining for proteins.

### Phage amplification and plaque assays

MG1655 cells were grown overnight in LB medium at 37 °C with shaking at 200 rpm, then diluted 1:100 into fresh LB medium, and grown to an OD_600_ of 0.4. Phages were added at an MOI of 0.1, and lysates were collected upon complete lysis. The lysates were clarified by centrifugation at 13,000 rpm for 25 min, filtered through a 0.22 μm sterile syringe filter and stored at 4 °C.

Wild-type and mutant DRT10-ncRNA constructs were cloned individually into the pBAD-LIC (Addgene #37501) vectors. The plasmids were transformed into *E. coli* MG1655. Transformants were grown overnight, diluted 1:100 in LB medium and cultured to an OD_600_ of 0.6. Gene expression was induced with 0.2% arabinose for 4 h. Then, 200 μL of induced culture was mixed with 10 mL of semi-solid agar containing the same concentrations of inducers and overlaid onto 10 cm agar plates. Serial tenfold dilutions of the phages were spotted onto the top agar, followed by overnight incubation at 37 °C.

### Cryo-EM sample preparation and data acquisition

For the DRT10-ncRNA complex, a 3.5 μL aliquot of purified sample at 0.87 mg/mL was applied to glow-discharged Quantifoil Au R1.2/1.3 holey carbon grids (300 mesh). The grids were blotted for 3 s at a blot force of 1 and a waiting time of 10 s at 4 °C and 100% humidity, then plunged into liquid ethane using a Vitrobot Mark IV (Thermo Fisher Scientific). Cryo-EM data for DRT10-ncRNA complexes were collected on a Titan Krios G4 microscope (Thermo Fisher Scientific) operated at 300 kV and equipped with a Falcon 4 direct electron detector. Data were acquired using EPU software (Thermo Fisher Scientific) at a magnification of 165,000×, corresponding to a calibrated pixel size of 0.725 Å, with a defocus range of -1.4 to -1.6 μm and a total dose of ∼50 e^-^/Å².

For the DRT10-ncRNA complex incubated with dNTPs, 0.5 mM dNTPs were incubated with the purified DRT10-ncRNA complex at 0.85 mg/mL in a buffer containing 25 mM Tris-HCl pH 7.5, 300 mM NaCl, 5 mM MgCl_2_ and 2 mM DTT. The mixture was incubated at room temperature for 5 min. An aliquot of 3.5 μL sample was then vitrified as described above, except that the grids were blotted for 2 s with a force of 0 and a waiting time of 5 s. Grid was transferred to Titan Krios G4 microscope and imaged with a calibrated pixel size of 0.731 Å.

### Cryo-EM image processing

Cryo-EM datasets of the DRT10-ncRNA complex prepared in the presence or absence of dNTPs were preprocessed in CryoSPARC using patch motion correction and patch CTF estimation (30). For the DRT10-ncRNA complex in the dNTP-free state, particles were picked by combining blob picking and Topaz picking (31). All picked particles were extracted with a box size of 360 pixels and classified with multiple rounds of 2D classification. The 546,789 particles with well-defined structural features were selected to perform ab-initio reconstruction and heterogeneous refinement. Particles contributing to the highest-quality reconstruction were further subjected to 3D classification and non-uniform refinement to obtain the final reconstruction at 2.59 Å. For the DRT10-ncRNA complex in the dNTP-bound and product-bound states, particles were picked as described above. After being extracted with a box size of 400 pixels, 496,846 particles were subjected to ab-initio reconstruction and heterogeneous refinement. Subsequent 3D classification and non-uniform refinement yielded the final reconstructions of dNTP-bound and product-bound states at overall resolutions of 2.81 and 3.44 Å, respectively.

### Model building and refinement

The initial model of the DRT10-ncRNA complex was predicted by AlphaFold3 (32). The predicted model was docked into the cryo-EM map using ChimeraX software (33). Subsequent manual adjustment and refinement were performed in Coot (34). After iterative docking and manual adjustment, the model was refined automatically using phenix.real_space_refine and validated with MolProbity in Phenix (35,36). The models in the dNTP-bound and product-bound states were modeled using the same procedure. Data collection and refinement statistics are provided in Table S1.

---

**Figure S1.**
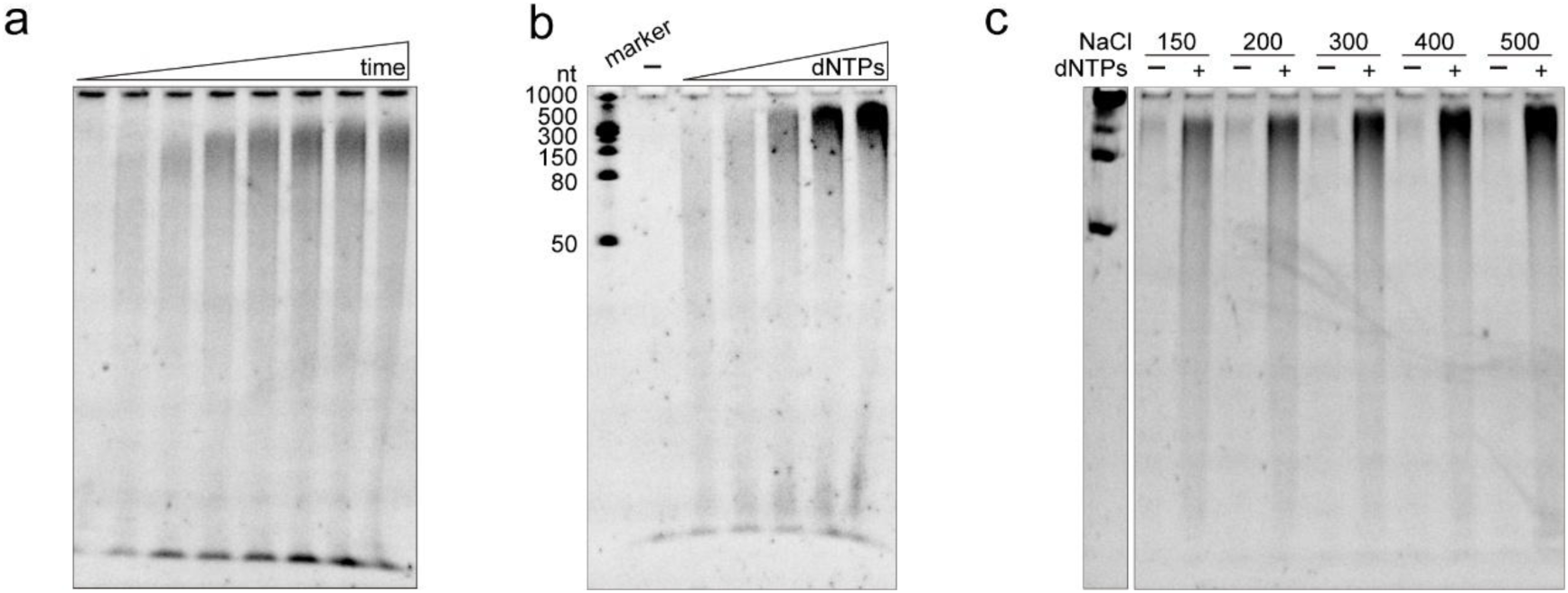
In vitro reverse transcription assay of DRT10-ncRNA complex. **a-c** In vitro reverse transcription assay of DRT10-ncRNA complex performed over a range of incubation times, dNTP concentrations and salt concentrations. The representative results are from three replicates.

**Figure S2.**
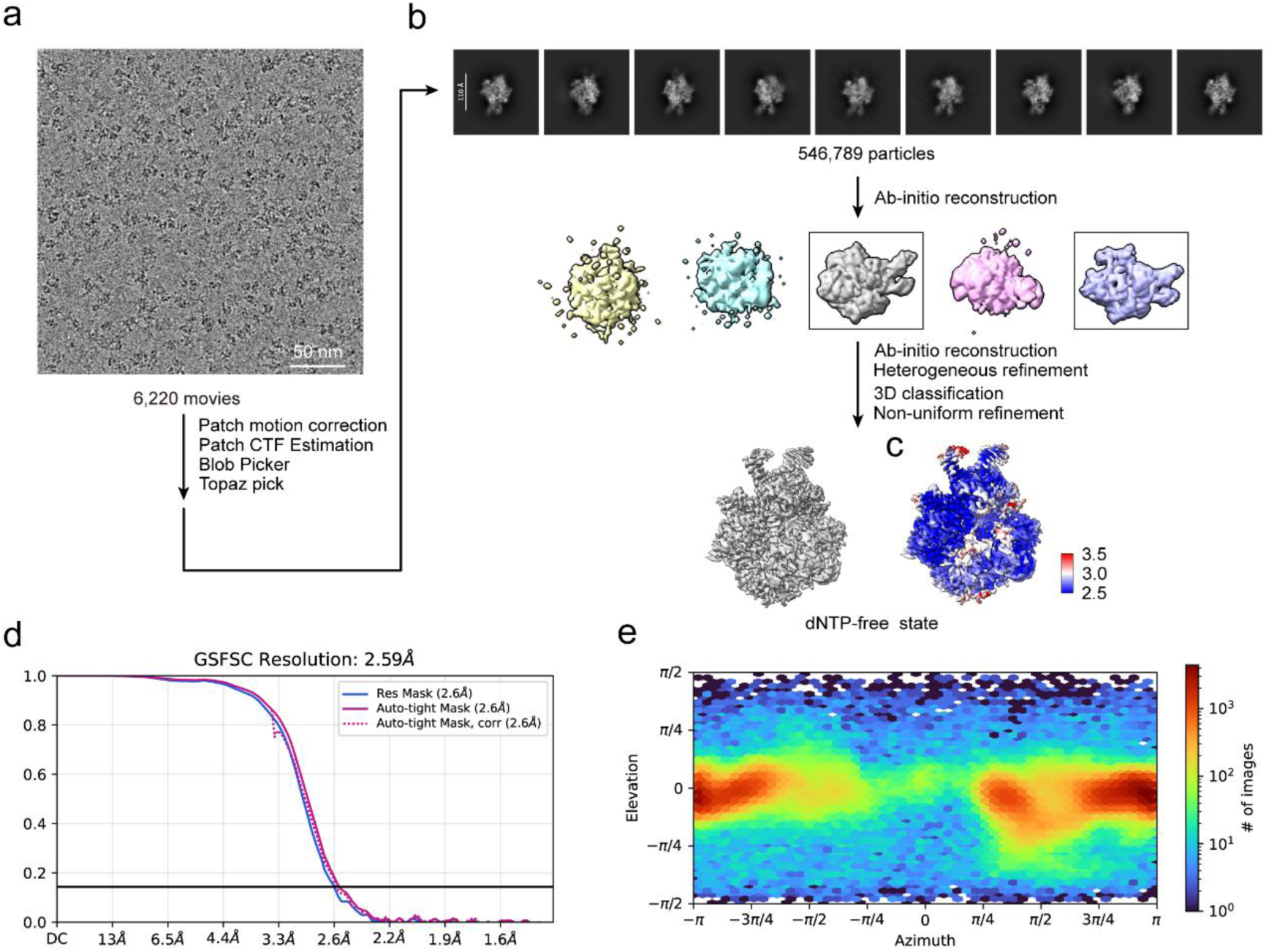
Cryo-EM data processing of DRT10-ncRNA complex in the dNTP-free state. **a** The data-processing workflow for DRT10-ncRNA complex in the dNTP-free state. **b** Images of representative 2D class average. **c** Local resolution map of DRT10-ncRNA complex in the dNTP-free state. **d** Fourier shell correlation (FSC) curve for the final reconstruction estimated at a cutoff of FSC = 0.143. **e** Viewing direction distribution of the reconstruction.

**Figure S3.**
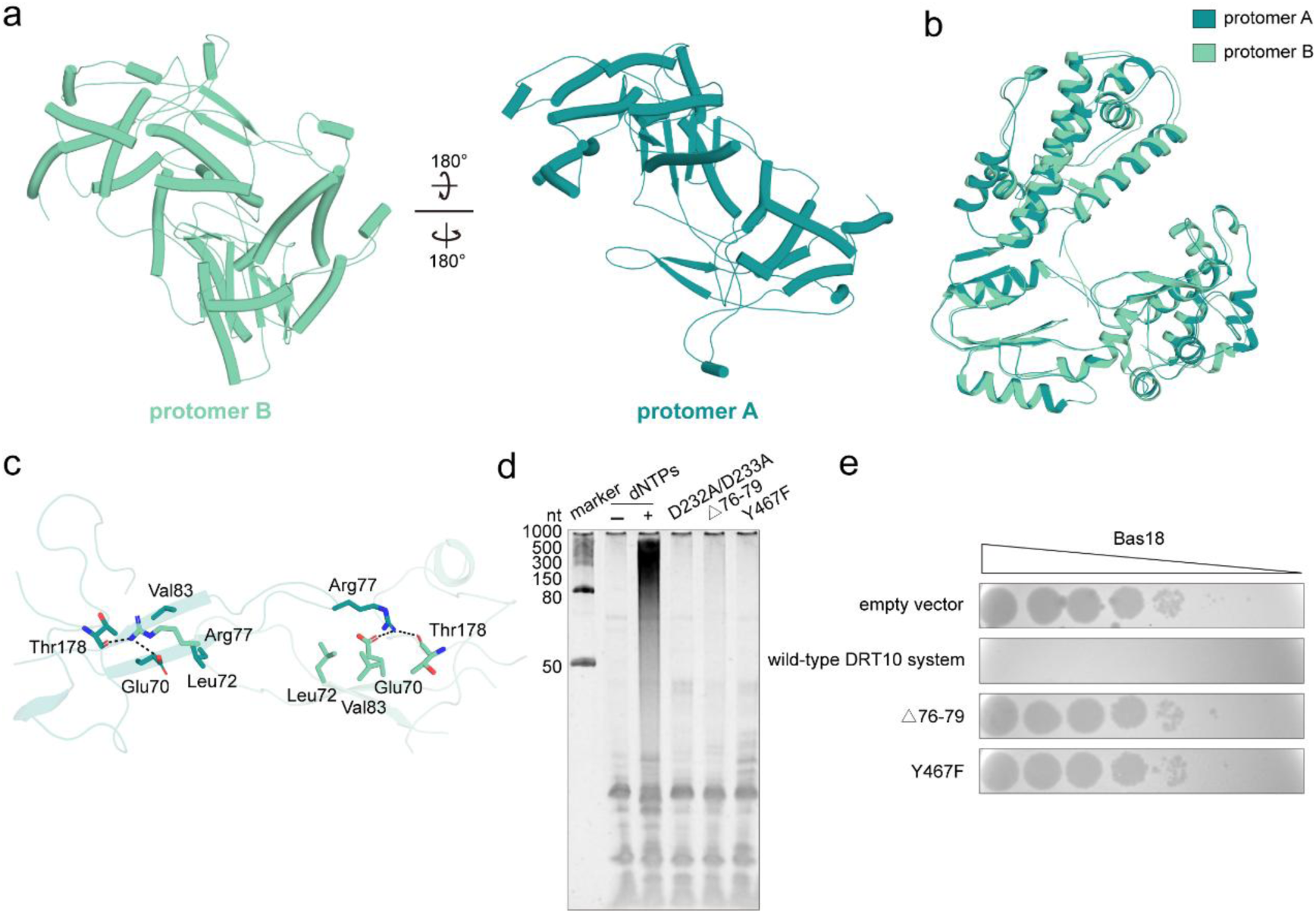
Interlocked β-hairpins are essential for DRT10 reverse-transcription activity and antiphage defense. **a** Atomic models of two DRT10 protomers in the dNTP-free state. **b** Structural alignment of two DRT10 protomers in the dNTP-free state. **c** Close-up view of the interactions formed by the interlocked β-hairpins. **d** In vitro reverse transcription assay of wild-type and indicated mutant DRT10-ncRNA complexes. A representative gel image from three experiments is shown. **e** Plaque formation by serially diluted phage Bas18 on MG1655 lawns carrying the indicated wild-type or mutant DRT10 constructs and the empty-vector control. Representative images are from three replicates.

**Figure S4.**
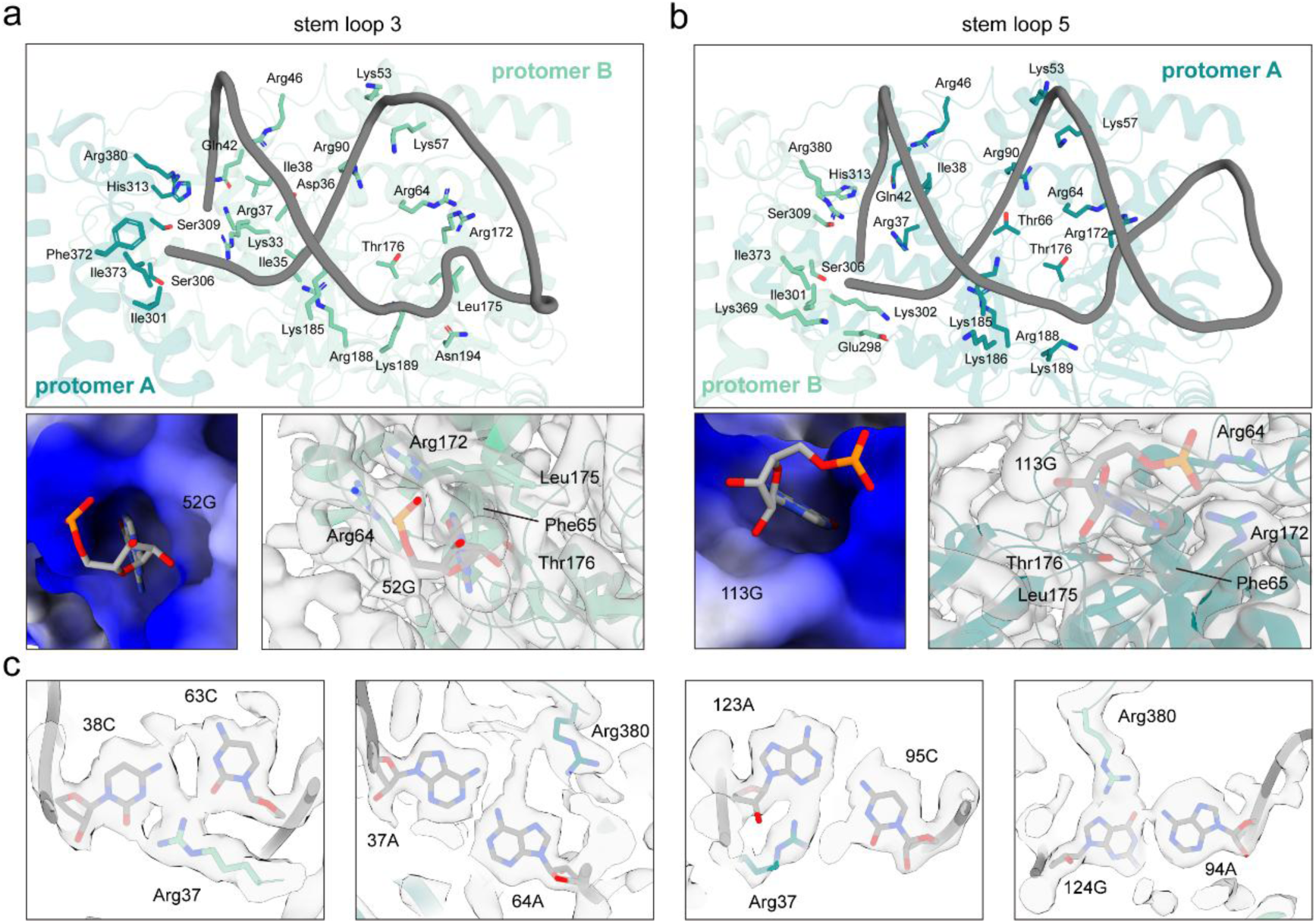
Extensive interactions of stem loop 3 and stem loop 5 with the DRT10 protomers. **a-b** Detailed interaction networks between stem loop 3 (a) or stem loop 5 (b) and the corresponding DRT10 protomer. The lower-left panels show the electrostatic surface potentials of the pockets accommodating 52G and 113G. The lower-right panels show the cryo-EM densities for 52G and 113G and their interacting residues. Key residues are shown as sticks. **c** Cryo-EM densities and atomic models of the noncanonical base-pairing interactions within stem loop 3 and stem loop 5.

**Figure S5.**
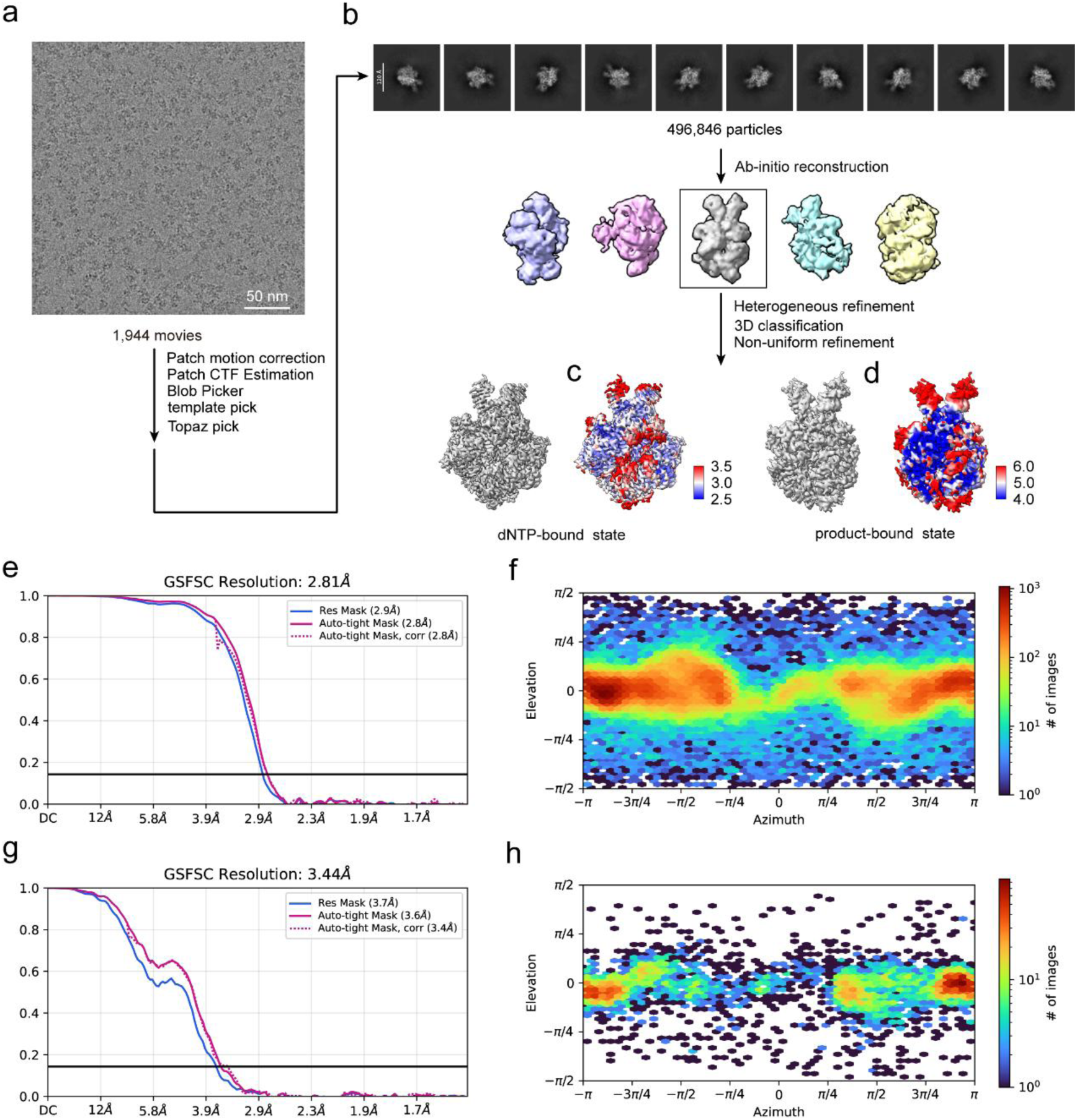
Cryo-EM data processing of DRT10-ncRNA complex in the dNTP-bound and product-bound states. **a** The data-processing workflow for DRT10-ncRNA complex incubated with dNTPs. **b** Representative 2D class averages. **c-d** Local-resolution maps of the final reconstructions in the dNTP-bound (**c**) and product-bound (**d**) states. **e** FSC curve for the final reconstruction of DRT10-ncRNA complex in the dNTP-bound state, with the overall resolution determined using the FSC = 0.143 criterion. **f** Viewing direction distribution of the reconstruction in the dNTP-bound state. **g** FSC curve for the final reconstruction of DRT10-ncRNA complex in the product-bound state. **h** Viewing direction distribution of the reconstruction in the product-bound state.

**Figure S6.**
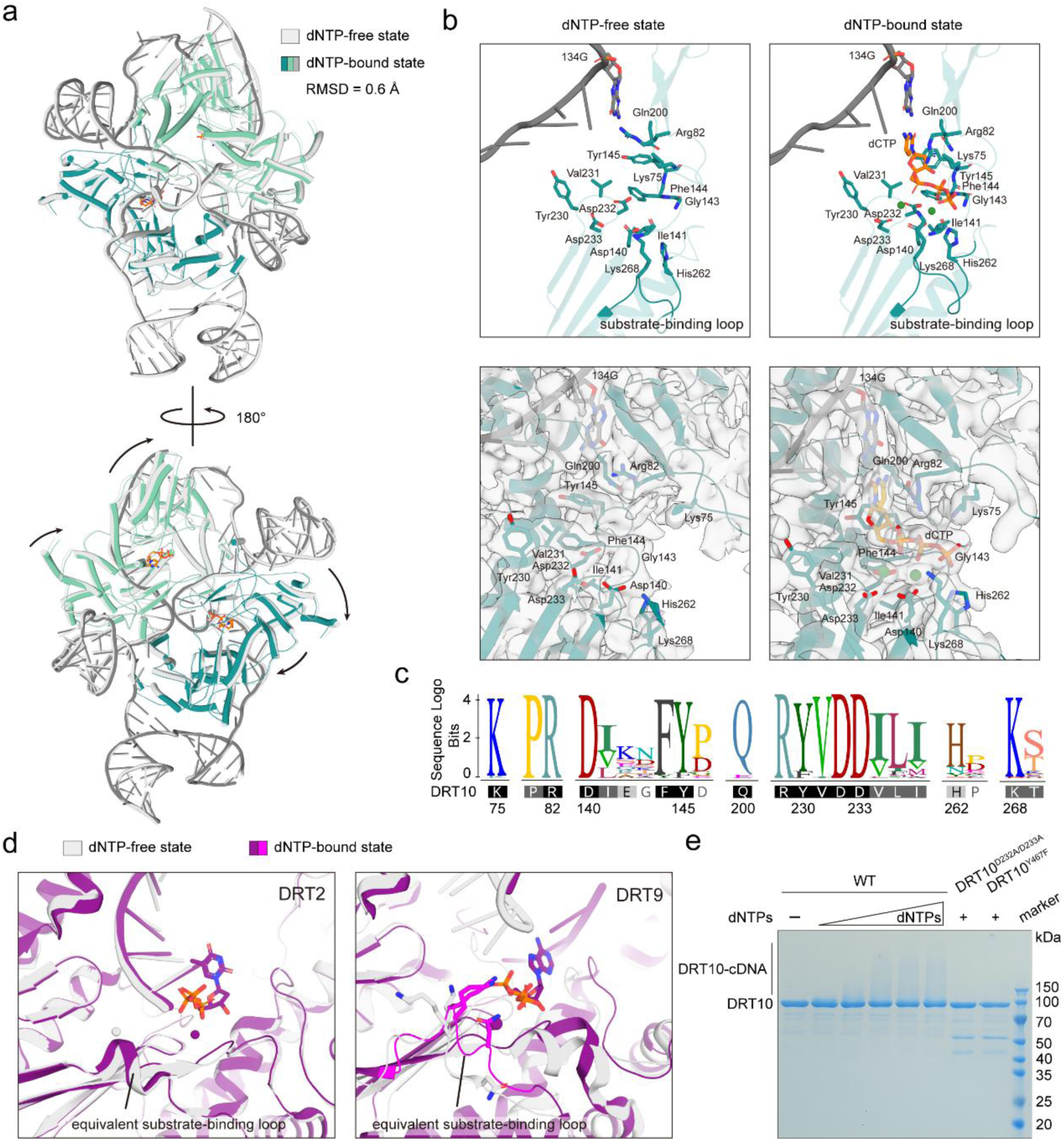
The substrate-binding loop facilitates dNTP engagement. **a** Structural alignment of DRT10-ncRNA complexes in the dNTP-free and dNTP-bound states. **b** Key residues involved in dNTP binding in the dNTP-free (left) and dNTP-bound (right) states. **c** Sequence logo showing the conservation of residues involved in dNTP binding among DRT10 homologs. **d** Structural comparison of the corresponding substrate-binding loops in the dNTP-free and dNTP-bound structures of DRT2 (PDB 9C0I and 9C0J) and DRT9 (PDB 9VKU and 9VMA). **e** In vitro reverse transcription assays of the wild-type DRT10-ncRNA complex and the indicated variants, analyzed by SDS-PAGE. Representative gel image is derived from three experiments.

**Figure S7.**
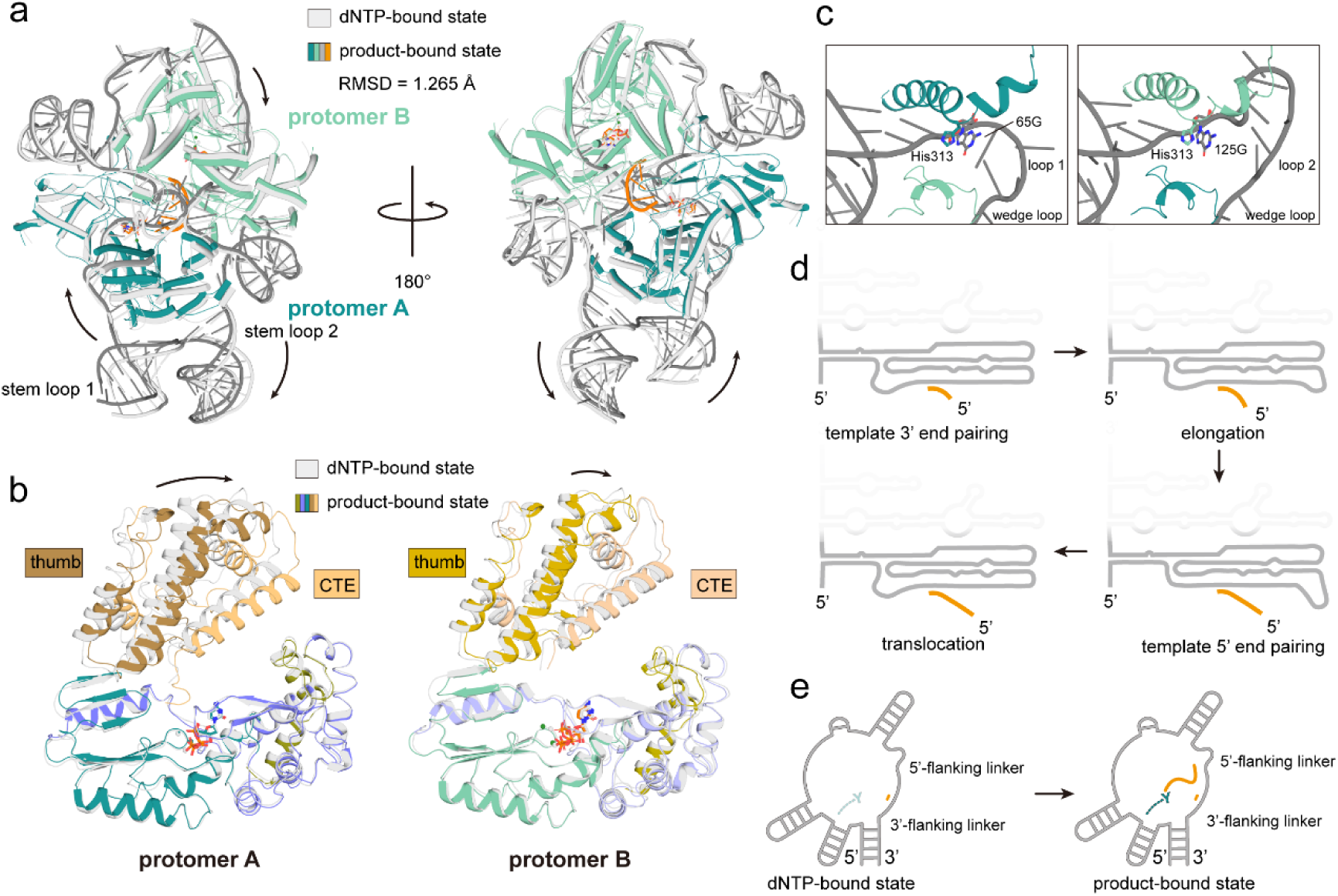
Structural comparison of DRT10-ncRNA complex in the dNTP-bound and product-bound states. **a** Structural alignment of DRT10-ncRNA complexes in the dNTP-bound and product-bound states. **b** Superposition of the corresponding DRT10 protomers in the dNTP-bound and product-bound states. **c** Close-up views of the 5’ ends of 5’-flanking linkers in loop 1 (left panel) and loop 2 (right panel). **d** Schematic representation of the RNA accordion model for telomerase reverse transcription. **e** Schematic representation of reverse transcription mechanism of the DRT10 protein.

**Table S1.** Cryo-EM data collection, refinement and validation statistics.

|  | DRT10 complex in<br>the dNTP-free state | DRT10 complex in<br>the dNTP-bound state | DRT10 complex in the<br>product-bound state |
| --- | --- | --- | --- |
| **Data collection and processing** |  |  |  |
| Magnification | 165,000 | 165,000 | 165,000 |
| Voltage (kV) | 300 | 300 | 300 |
| Electron exposure (e<sup>-</sup>/Å<sup>2</sup>) | 50.30 | 50 | 50 |
| Defocus range (μm) | -1.4 to -1.6 | -1.4 to -1.6 | -1.4 to -1.6 |
| Pixel size (Å) | 0.725 | 0.731 | 0.731 |
| Symmetry imposed | *C1* | *C1* | *C1* |
| Initial particle images (no.) | 546,789 | 496,846 | 496,846 |
| Final particle images (no.) | 336,153 | 115,750 | 5,774 |
| Map resolution (Å) | 2.59 | 2.81 | 3.44 |
| FSC threshold | 0.143 | 0.143 | 0.143 |
| Map resolution range (Å) | 2.5-3.5 | 2.5-3.5 | 4.0-6.0 |
| **Refinement** |  |  |  |
| Initial model used (PDB code) | AlphaFold3 | AlphaFold3 | AlphaFold3 |
| Model resolution (Å) | 2.8 | 3.0 | 3.7 |
| FSC threshold | 0.5 | 0.5 | 0.5 |
| Model resolution range (Å) | 2.5-2.8 | 2.7-2.9 | 3.3-3.7 |
| Map sharpening *B* factor (Å<sup>2</sup>) | -97.9 | -70.6 | -17.7 |
| Model composition |  |  |  |
| Non-hydrogen atoms | 10696 | 10709 | 10809 |
| Protein residues | 934 | 929 | 931 |
| Nucleotides | 140 | 140 | 144 |
| Ligands | 0 | 6 | 6 |
| *B* factors (Å<sup>2</sup>) |  |  |  |
| Protein | 35.06 | 60.40 | 90.86 |
| Nucleotide | 67.78 | 93.93 | 138.96 |
| Ligand | - | 47.03 | 84.77 |
| R.m.s. deviations |  |  |  |
| Bond lengths (Å) | 0.005 | 0.004 | 0.003 |
| Bond angles (°) | 0.874 | 0.746 | 0.643 |
| Validation |  |  |  |
| MolProbity score | 1.83 | 1.33 | 1.72 |
| Clashscore | 6.00 | 3.80 | 6.85 |
| Poor rotamers (%) | 1.87 | 0.59 | 0.23 |
| Ramachandran plot |  |  |  |
| Favored (%) | 95.81 | 97.08 | 95.15 |
| Allowed (%) | 4.09 | 2.92 | 4.85 |
| Disallowed (%) | 0.1 | 0 | 0 |

## Notes

### Competing Interest Statement

The authors have declared no competing interest.

